# A Generative AI Framework to Predict Cardiomyocyte Contraction Function from Single Static Images

**DOI:** 10.64898/2026.04.22.720172

**Authors:** Andrew Kowalczewski, Chenyan Wang, Xinrui Wang, Huaxiao Yang, Zhao Qin, Zhen Ma

**Affiliations:** Department of Biomedical & Chemical Engineering, Syracuse University, Syracuse NY, USA; BioInspired Syracuse Institute for Materials and Living Systems, Syracuse University, Syracuse NY, USA; Department of Pharmacology, SUNY Upstate Medical University, Syracuse, NY, USA; Department of Biomedical Engineering, University of North Texas, Denton, TX; Department of Civil & Environmental Engineering, Syracuse University, Syracuse NY, USA

**Keywords:** Artificial Intelligence (AI), Deep Learning, Cardiomyocytes, Human Induced Pluripotent Stem Cells (hiPSCs)

## Abstract

Understanding how cardiomyocyte structure governs contractile function is fundamental to cardiac biology and disease modeling, yet current approaches rely on time-resolved imaging and computationally intensive analysis. Here, we present a generative artificial intelligence (AI) framework that directly predicts contractile behavior of human induced pluripotent stem cell-derived cardiomyocytes (hiPSC-CMs) from single static images. Our approach integrates a U-Net–based generator with a patch-based generative adversarial network (GAN) discriminator to translate morphological and sarcomere structural features into pixel-resolved contraction heatmaps. This U-Net-GAN model achieved high predictive accuracy, with structural similarity index (SSIM) values up to 0.84 using combined morphological and structural inputs. To further enhance performance and generalizability, we incorporated synthetic cell–function pairs generated via a generative AI StyleGAN2 framework, improving prediction accuracy and perceptual similarity. Importantly, region-specific and whole-cell analyses revealed that AI predictions capture biologically meaningful structure–function relationships, with sarcomere organization strongly associated with both contractile output and prediction fidelity. Reconstruction error emerged as an interpretable metric reflecting localized inefficiencies in sarcomere-to-contraction coupling. Together, this framework establishes a scalable and interpretable strategy for inferring cardiomyocyte function from static morphology, eliminating the need for time-lapse imaging. More broadly, this work positions generative AI as a powerful tool for bridging cellular structure and function, enabling high-throughput functional phenotyping and advancing in vitro cardiac modeling.

## INTRODUCTION

Cardiomyocytes are highly specialized cells whose structure is tightly coupled to their function. The contractile apparatus of these cells is organized into striated, repeating sarcomeres, which are the fundamental units of active contraction^1–3^. At the cellular level, sarcomere length, alignment, and periodicity strongly correlate with contractile force generation^4–6^. Proper sarcomere organization is therefore essential for coordinated force generation and efficient contraction. This well-established structure-function relationship that structural features of the sarcomere network may serve as predictive indicators of cardiomyocyte contractile performance.

Human induced pluripotent stem cell-derived cardiomyocytes (hiPSC-CMs) offer a human-relevant platform for investigating the relationship between morphological and functional phenotypes in vitro^7,8^. Traditional high-content imaging assays, however, often require multiple staining steps and computationally intensive analyses to characterize structure and function of hiPSC-CMs^9–13^. Recent advances in deep learning have transformed biomedical image analysis, enabling extraction of complex spatial and functional information directly from microscopy images. For examples, convolutional neural networks (CNNs) can infer fluorescence signals, such as DAPI labeling, directly from brightfield images, a strategy commonly referred to as “virtual staining”^14^. Virtual staining involves mapping grayscale intensity information into multi-channel (RGB) representations that approximate fluorescent labeling. However, traditional CNN-based translation methods often produce blurred outputs that lack high-frequency detail and structural fidelity, partly due to pixel-wise averaging effects during reconstruction. Generative adversarial networks (GANs) have substantially improved this approach by incorporating an adversarial loss that encourages sharper and more realistic image synthesis, thus producing high-fidelity virtual stains that closely approximate experimentally acquired fluorescent images^15–17^. Recently, GAN-based virtual staining has been successfully applied to complex 3D cardioids, achieving high accuracy in reproducing cell-specific patterns^18^. Collectively, these studies demonstrate the capacity of deep learning to bridge morphological imaging and molecular characterization.

Despite these advances, no existing AI framework has quantitatively translated static cell morphology and sarcomere organization into direct predictions of contractile performance in hiPSC-CMs. To address this gap, we developed a traditional deep learning U-Net autoencoder and generative AI framework based on a U-Net architecture coupled with a patch-based GAN discriminator called a pix2pix model to predict cardiomyocyte contraction directly from single static microscopy images. To further enhance predictive performance, we augmented the training dataset with synthetic hiPSC-CMs generated using a StyleGAN2 framework, which produces paired synthetic cell images and corresponding contraction heatmaps. To improve biological interpretability, we systematically correlated traditional measurements of cell morphology, sarcomere structure, and contractile function with AI-derived metrics (Figure 1). Our analyses revealed that AI model integrates both morphological and structural features when inferring contractile behavior. This approach enables rapid prediction of single-cell contractile function from a single snapshot, eliminating the need for time-lapse imaging and extensive post-processing. By directly linking cellular structure to function, our framework establishes an interpretable and scalable strategy for high-throughput assessment of hiPSC-CM contractility.

**Figure 1.**
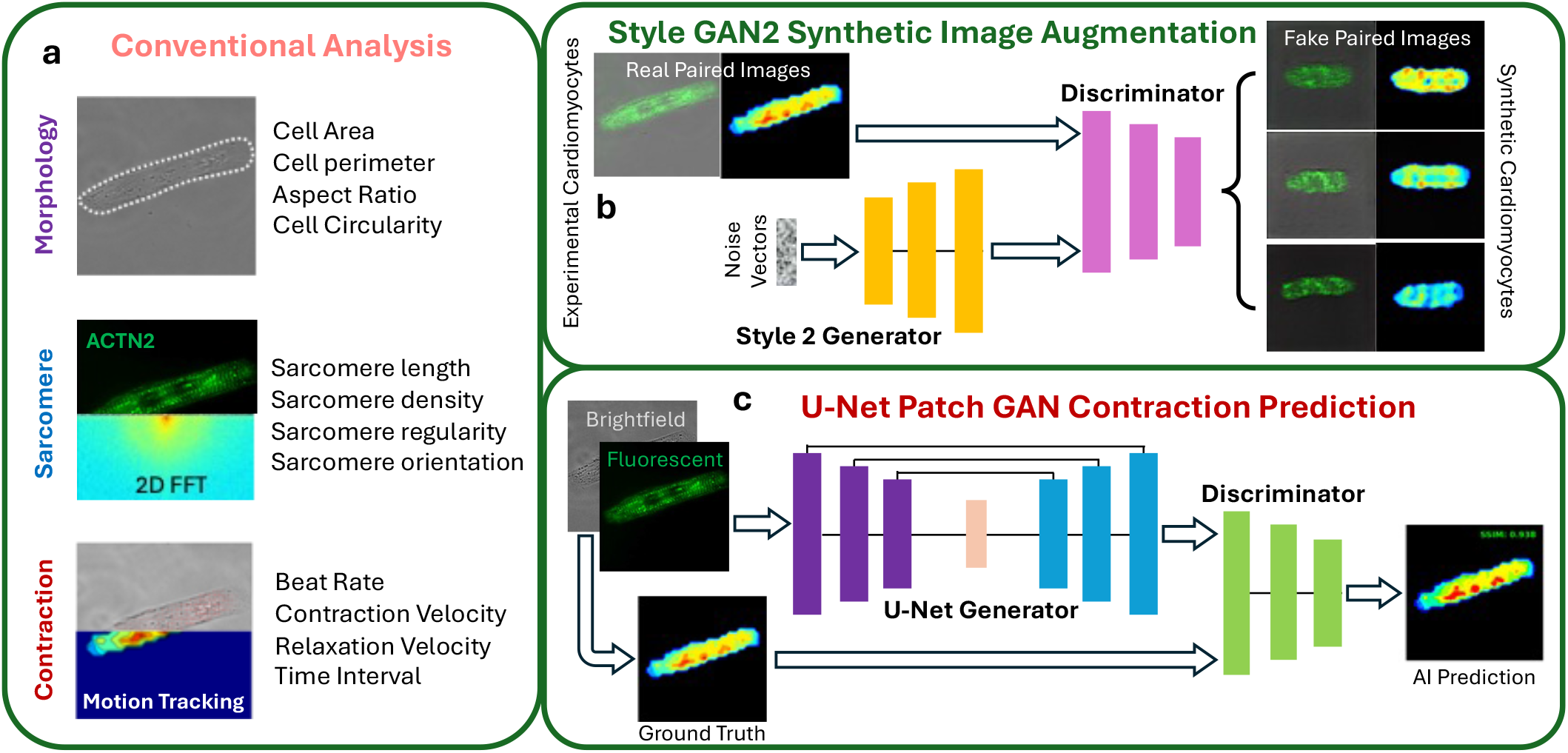
Overview of the analytical framework. (a) Conventional analysis of cell morphology, sarcomere structure, and contractile function. (b) StyleGAN2 framework for generating synthetic hiPSC-CMs and corresponding contraction heatmaps. (c) U-Net–GAN model for predicting contraction heatmaps from static images.

## RESULTS

### Predicting Contraction Function from Single Static Images

hiPSC-CMs were differentiated and purified from a CRISPR-engineered ACTN2-GFP reporter line, allowing direct live-cell imaging of sarcomere structures. Following differentiation and purification, cells were micropatterned using a microcontact printing approach (Supplemental Figure 1a). Fibronectin islands were first printed onto glass substrates (Supplemental Figure 1b), after which hiPSC-CMs were seeded and cultured on these patterned surfaces (Supplemental Figure 1c). Although micropatterning was used to guide cell organization, we did not selectively choose cells with idealized geometries. Instead, we imaged a large and morphologically diverse population of hiPSC-CMs to train and evaluate the generalizability of our generative AI framework, as reflected in the variability observed in the testing dataset (Supplemental Figure 1d).

Cells were imaged using brightfield microscopy to capture cell morphology and contractile motion (Movie S1) and fluorescence microscopy to visualize sarcomere structure. Contractile motion was quantified from the brightfield videos using a robust, open-source block-matching motion analysis algorithm^9^. In total, 296 hiPSC-CMs were collected with paired image datasets consisting of: (1) brightfield morphology images, (2) fluorescent sarcomere images, (3) merged brightfield/fluorescent overlays, and (4) corresponding contractile heatmaps derived from block-matching motion tracking of brightfield videos. These heatmaps provided pixel-level ground-truth measurement of contractile displacement intensity for each cell. The dataset was partitioned into a training set (n = 266) and an independent testing set (n = 30) to train and evaluate the AI model for predicting contractile behavior from static single-cell images.

To investigate whether generative AI could translate a single static snapshot into predicted contractile function, we implemented a U-Net–based generative adversarial network (U-Net-GAN), adapted from pix2pix-style image-to-image translation frameworks^17–19^. In this architecture, the U-Net autoencoder serves as the generator, while a 70×70 patch-GAN discriminator supervises the generator to enhance structural fidelity and local realism (Figure 2a). Rather than evaluating entire images globally, the patch-based discriminator enforces local consistency, encouraging reconstruction of fine spatial features and subcellular textures that are critical for accurate functional mapping^20,21^.

**Figure 2.**
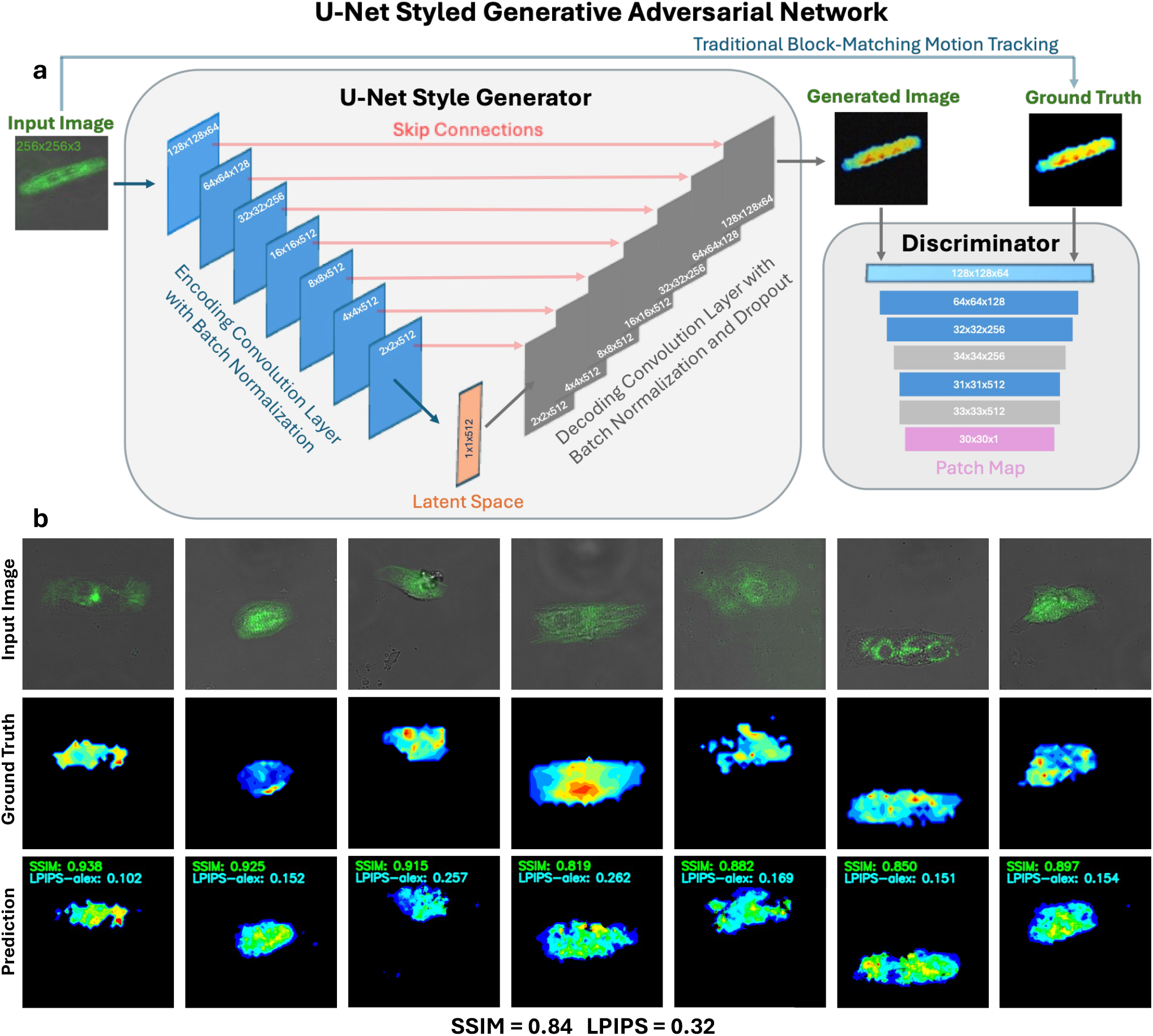
Translation from static cell images to contraction heatmaps. (a) Schematic of the U-Net–GAN generative AI framework used to predict contraction heatmaps from single static images of hiPSC-CMs. (b) Representative examples from the testing dataset showing paired inputs and outputs: an overlay image of cell morphology (brightfield) and sarcomere structure (fluorescence), the corresponding ground-truth contraction heatmap computed using a block-matching algorithm, and the AI-predicted contraction heatmap generated by the U-Net–GAN model.

The model was trained using input hiPSC-CM images (brightfield morphology, fluorescent sarcomere images, or their overlays) as inputs and corresponding ground-truth contraction heatmaps as outputs. On the held-out testing dataset, the U-Net-GAN achieved a structural similarity index (SSIM) of 0.80 when translating brightfield images into contraction heatmaps (Supplemental Figure 2a). Using fluorescent sarcomere images as input improved predictive performance, yielding an SSIM of 0.82 (Supplemental Figure 2b), even though the ground truth heatmaps were computed from brightfield videos. To further enhance predictive accuracy, we integrated morphological and structural information by overlaying brightfield and fluorescence images as a combined input. This refinement increased model performance to an SSIM of 0.84 on the testing dataset (Figure 2b). The predicted contraction heatmaps exhibited sharper boundaries and improved spatial localization of contractile regions compared to single-modality inputs.

### Synthetic Cardiomyocytes to Improve Model Performance

To further improve the performance and generalizability of our contraction prediction model, we investigated whether synthetic hiPSC-CMs could be generated to replicate key features of experimentally derived cells and used to augment U-Net–GAN training. We adopted a StyleGAN2 framework, which has demonstrated strong performance in generative image tasks and transfer learning with limited datasets^22,23^. StyleGAN2 constructs a latent space via a mapping network that transforms a random Gaussian noise vector into an intermediate latent representation, which is then injected into each layer of the generator (Figure 3a). This hierarchical architecture enables progressive feature synthesis, where early layers capture coarse attributes such as cell shape and size, while deeper layers refine fine-scale features including sarcomere organization and contraction patterns. A total of 81 high-quality synthetic cell–heatmap pairs were generated (Supplemental Figure 3), curated, and incorporated into the training dataset alongside 296 experimentally acquired hiPSC-CMs, resulting in an augmented dataset of 377 samples (Figure 3b). The inclusion of synthetic samples increased diversity in cell morphology, sarcomere organization, and fluorescence intensity distributions, thereby enriching the training space. Incorporation of synthetic hiPSC-CMs improved model performance (SSIM increased from 0.84 to 0.86; LPIPS decreased from 0.32 to 0.26) on the same held-out testing set.

**Figure 3.**
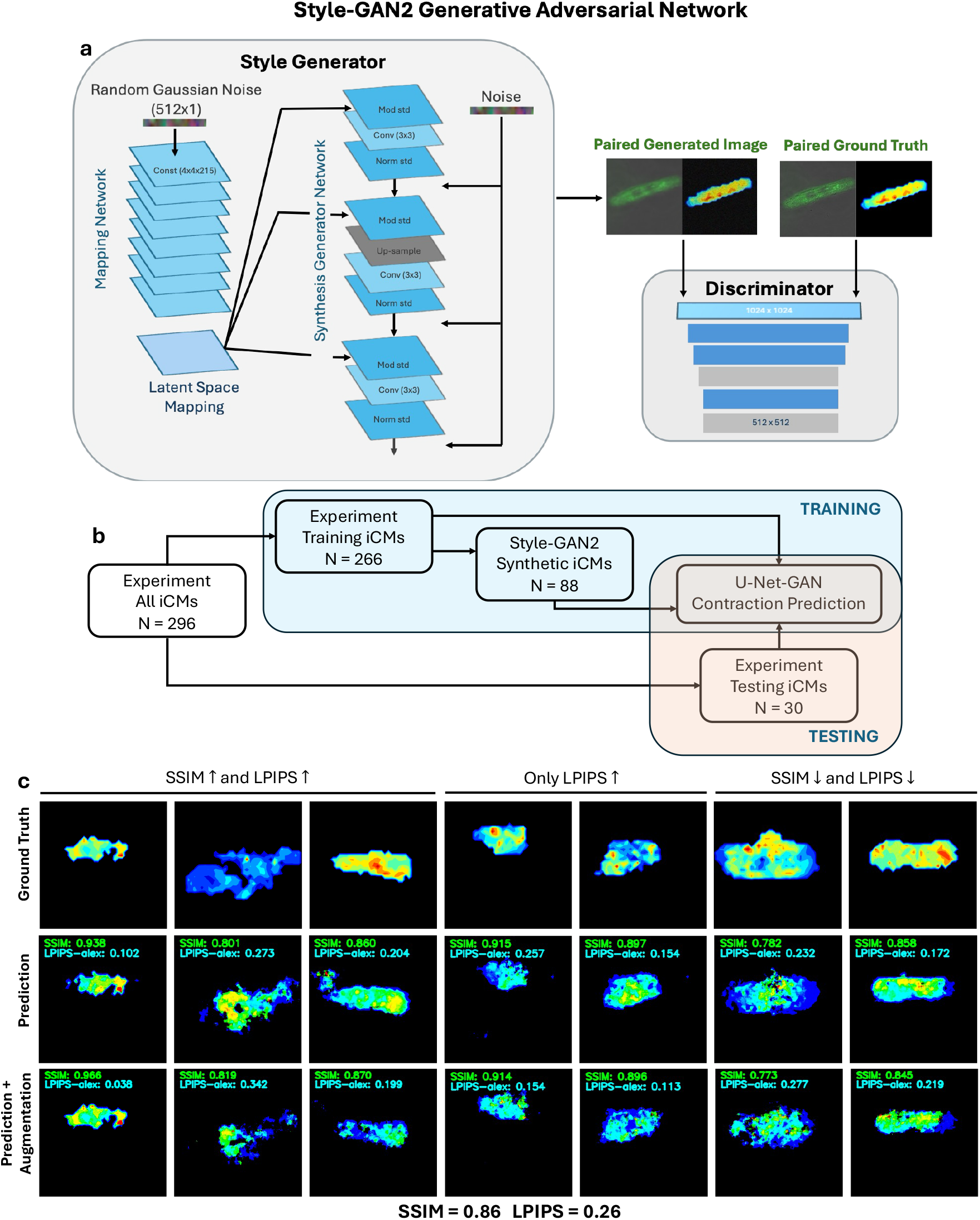
Generation of synthetic hiPSC-CMs to improve contraction prediction. (a) Schematic of the Style-GAN2 generative framework used to generate paired synthetic hiPSC-CM images and corresponding contraction heatmaps. (b) Real (experimentally acquired) and synthetic hiPSC-CMs were combined to train the U-Net–GAN model, which was evaluated on the same held-out testing dataset. (c) Representative examples from the testing set showing ground-truth contraction heatmaps and AI-predicted heatmaps generated with and without augmentation by synthetic hiPSC-CMs. Overall, the augmentation improves SSIM and LPIPS metrics.

### AI-Informed Morphology-Structure-Function Relationship

To improve interpretability of the AI model’s predictions, we analyzed region-specific relationships between sarcomere organization, contraction intensity, and reconstruction error within individual testing hiPSC-CMs. For each cell, contraction regions of interest (ROIs) were defined as areas of high versus low mean contraction intensity based on ground-truth contraction heatmaps. Error ROIs were defined as areas of high versus low reconstruction error, calculated from the pixel-wise difference between ground-truth and AI-predicted heatmaps (Figure 4a). Within these ROIs, sarcomere structural features, including sarcomere length, regularity, orientation, and density, were quantified using an automated batch-analysis pipeline based on two-dimensional Fast Fourier Transform (2D FFT), a well-established method for assessing sarcomere periodicity and organization in cardiomyocytes^24^. Analysis of ground-truth contraction heatmaps revealed that high-contractility regions exhibited significantly longer sarcomere length and higher sarcomere density compared to low-contractility regions (Figure 4b). Notably, when stratifying regions by AI reconstruction error, low-error regions (i.e., areas of higher predictive accuracy) demonstrated similarly increased sarcomere length and density relative to high-error regions (Figure 4c). This consistent structural trend across both contraction intensity and prediction accuracy suggests that localized sarcomere disorganization contributes to diminished contractile performance and reduced model fidelity. In this context, reconstruction error serves as a functional readout of structural inefficiency, revealing localized breakdowns in the translation of sarcomere organization into cardiac contraction.

**Figure 4.**
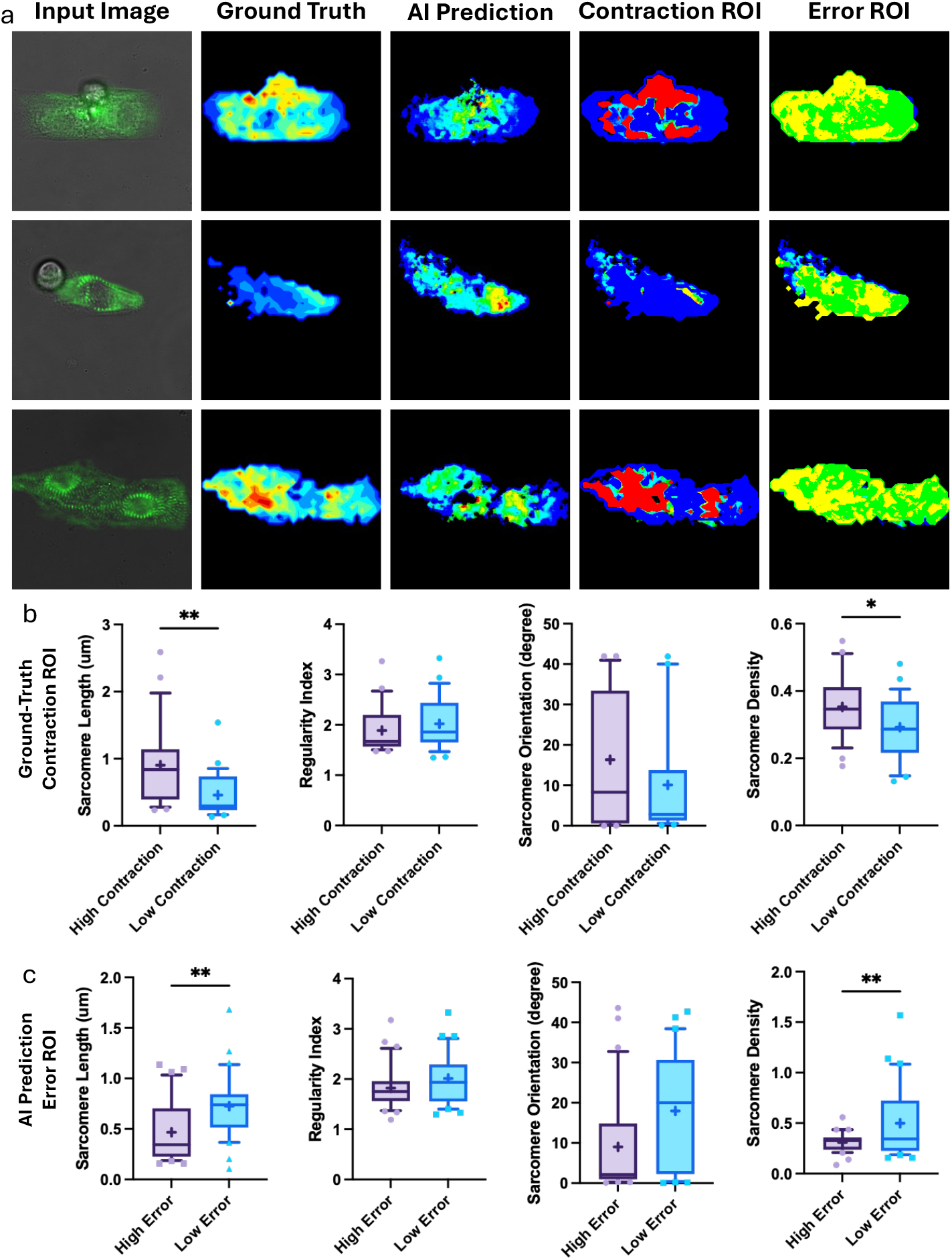
Region-specific structure-function relationship. (a) For each testing hiPSC-CM, regions of interest (ROIs) were segmented based on contraction intensity (ground truth) and reconstruction error (AI prediction) to enable region-specific comparison of sarcomere structural features. (b) Comparison between high- and low-contraction ROIs shows that high-contraction regions exhibit significantly greater sarcomere length and higher sarcomere density. (c) Comparison between high- and low-error ROIs shows that low-error regions (i.e., higher prediction accuracy) exhibit greater sarcomere length and higher sarcomere density.

To further examine global determinants of AI predictivity, we analyzed whole-cell features and their relationships to AI-related performance metrics, including SSIM, learned perceptual image patch similarity (LPIPS), high-to-low contraction area ratio (H/L contraction), and high-to-low error area ratio (H/L error). For each testing hiPSC-CM, we quantified cell morphology (area, perimeter, circularity, aspect ratio), sarcomere organization (length, regularity, orientation, density), and beating behaviors (beat rate, maximal contraction velocity, maximal relaxation velocity, and contraction–relaxation time interval). Spearman correlation analysis revealed strong intra-modality correlations (Figure 5a). For example, cell area and perimeter were highly correlated within cell morphology; contraction and relaxation velocities were tightly correlated within beating dynamics; and sarcomere length and regularity were correlated within structural features.

**Figure 5.**
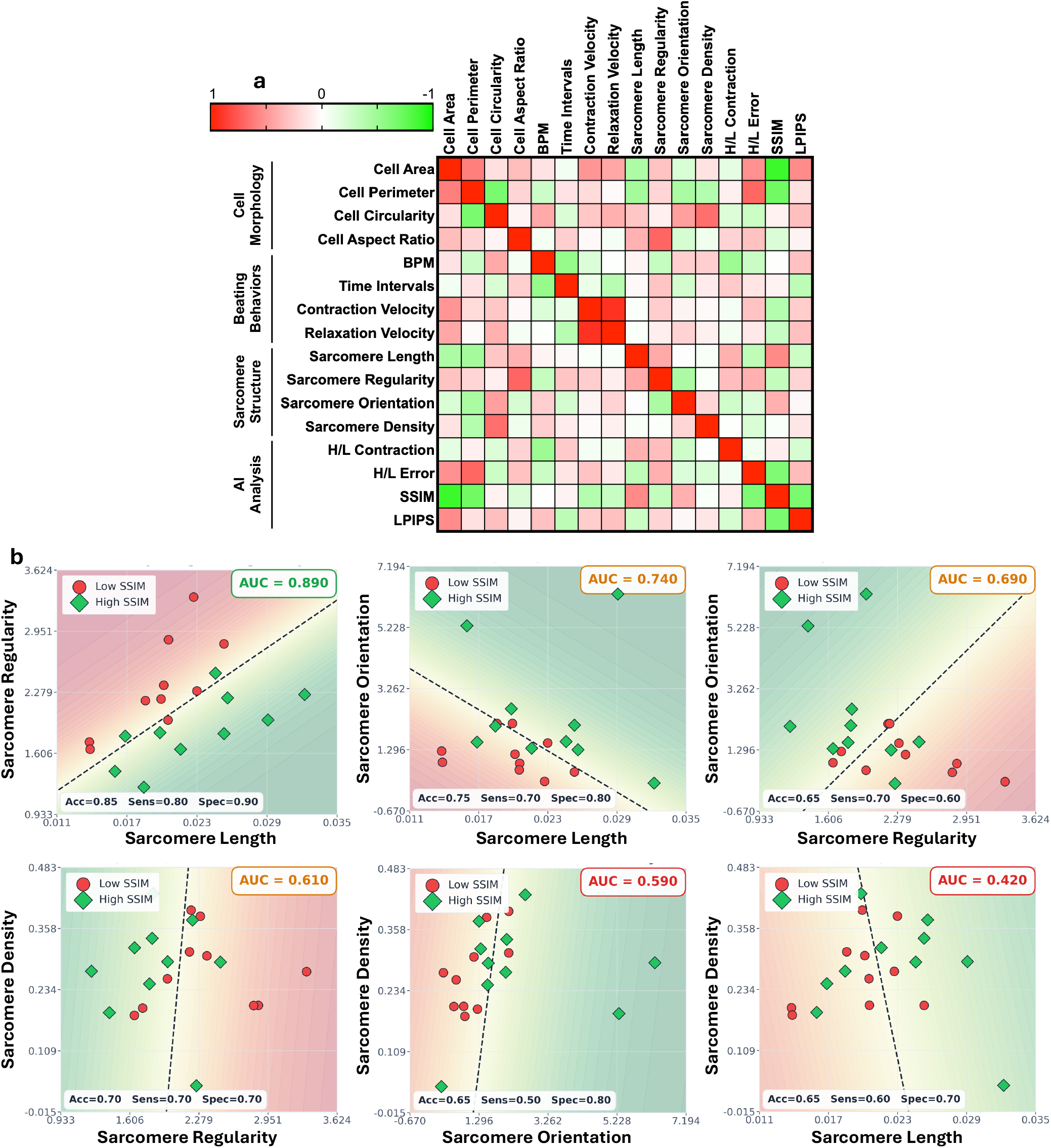
Biological Interpretation of AI predictivity metrics. (a) Correlation matrix across four cell morphology features, four sarcomere structural features, four contractile function features, and four AI predictivity metrics (red: positive correlation; green: negative correlation). (b) Logistic regression model was used to classify cells into high- and low-predictability groups based on paired sarcomere features. Decision boundaries are shown as color-filled contour maps in the two-dimensional feature space, with background probabilities (green: high possibility of predicting a cell with a high SSIM; red: low possibility).

Across modalities, cell aspect ratio positively correlated with sarcomere regularity (Supplemental Figure 4a), while cell circularity positively correlated with sarcomere density (Supplemental Figure 4b), suggesting anisotropic cell shape influence sarcomere organization. Functionally, the H/L contraction ratio was negatively correlated with beat rate (Supplemental Figure 4c) but positively correlated with sarcomere regularity (Supplemental Figure 4d). This indicates that improved sarcomere organization supports stronger contraction but may associate with reduced spontaneous beating frequency. Regarding AI predictivity, SSIM was negatively correlated with cell area and perimeter (Supplemental Figure 4e & 4f), suggesting that smaller hiPSC-CMs yielded more accurate predictions. As expected, there are strong correlation among these AI-related metrics, SSIM, H/L error and LPIPS (Supplemental Figure 4g – 4i).

Given the weak correlations between individual sarcomere features and AI predictivity metrics, we next examined whether combinations of structural features could jointly classify hiPSC-CMs into high-versus low-predictability groups (Figure 5b). Cells were dichotomized based on SSIM into high-SSIM (SSIM ≥ median = 0.842) and low-SSIM groups. We then performed logistic regression using all six pairwise combinations of four sarcomere features (length, regularity, orientation, and density). Model performance was evaluated using leave-one-out cross-validation (LOO-CV) with the area under the receiver operating characteristic curve (AUC). Among all feature pairs, the combination of sarcomere length and regularity achieved the highest classification performance (AUC = 0.890), followed by the length–orientation pair (AUC = 0.740). The resulting logistic decision boundary showed clear separation between high- and low-SSIM cells in the sarcomere length–regularity feature space, indicating that these features provide complementary structural information governing AI predictability.

## DISCUSSION

In this study, we developed a U-Net–GAN framework capable of predicting the contractile behavior of individual hiPSC-CMs from a single static image. This approach bypasses conventional motion-tracking pipelines and provides a scalable and generalizable strategy for inferring functional output directly from morphological and structural features. To further enhance model performance and generalizability, we implemented a StyleGAN2 framework to generate synthetic hiPSC-CMs paired with corresponding contraction heatmaps for data augmentation. In parallel, our analytical framework integrates automated cell morphology analysis and 2D FFT–based sarcomere quantification at both global and region-specific levels. Together, this Python-based contractile analysis toolkit provides a rapid, automated, and objective platform for cardiomyocyte functional assessment, substantially reducing experimental burden, human bias, and computational overhead.

### AI Applications in hiPSC-based cardiac models

The first wave of integrating machine learning (ML) and deep learning (DL) into in vitro iPSC-based cardiac models primarily focused on drug cardiotoxicity prediction and phenotypic classification^25–28^. Early studies applied ML models and CNNs to analyze motion waveforms, calcium transients, and electrophysiological recordings from hiPSC-CMs exposed to pharmacological perturbations^29–31^. These approaches enabled automated detection of arrhythmias, beat irregularities, and dose-dependent changes in contractility across diverse patient-specific iPSC lines. More recently, attention has expanded toward AI-assisted quality control of hiPSC-CM differentiation and maturation^32–34^. For instance, GAN–based models have been used in 3D cardiac organoids for automated cell-type annotation^18^ and in 2D hiPSC-CMs for differentiation and maturation optimization^35^.

Beyond classification and quality assessment, AI is increasingly functioning as a biological translation engine by bridging modalities, scales, and developmental stages in cardiac systems. For example, a long short-term memory (LSTM) multitask network has been developed to translate immature hiPSC-CM action potential waveforms into predicted adult-like electrophysiological profiles^36^. Similarly, physics-informed deep learning frameworks based on U-Net architectures with attention modules have reconstructed intracellular action potentials from extracellular field potential recordings^37^. Within this broader paradigm, image-to-function translation represents a biologically meaningful extension of generative modeling. Rather than merely enhancing image quality or performing virtual staining, our framework translates static morphological and sarcomere organization into dynamic mechanical output.

These advances suggest a unifying trajectory in which AI models will learn the continuous coupling between structure, function, and time. Cross-modal translation frameworks may ultimately enable integrated digital representations of cardiomyocytes, where morphology, electrophysiology, and mechanics are computationally linked, providing a scalable foundation for disease modeling, maturation assessment, and therapeutic optimization.

### Unified computational metrics for sarcomere-contraction coupling

Although the concept of sarcomere–contraction coupling is fundamental to cardiac biology, no established quantitative metric currently exists to directly measure the efficiency with which sarcomere organization translates into contractile performance at the single-cell level. Existing analyses typically quantify structure (e.g., sarcomere length, alignment)^32^ and function (e.g., displacement, calcium transient)^9,38^ independently, without an integrated framework to assess how effectively structural order is converted into mechanical output. Our AI-based model provides a data-driven approximation of this structure–function coupling. The strong predictive performance of the network supports the central premise that structural features embedded in a single image, including sarcomere periodicity, alignment, and cytoskeletal integrity, encode sufficient information to approximate contractile behaviors^39,40^. Importantly, we demonstrate that reconstruction error between AI-predicted and ground-truth contraction heatmaps is not merely a technical artifact but a biologically interpretable metric. Regions with high reconstruction error were significantly associated with sarcomere disorganization, indicating localized inefficiency in translating structural integrity into coordinated contraction. In addition, SSIM showed strong correlations with cell area and perimeter and could be effectively classified using paired sarcomere features, particularly length and regularity. Together, these findings suggest that AI-derived metrics can serve as computational proxies for morphology–structure–function coupling efficiency, capturing this relationship within a unified quantitative framework. This approach establishes a scalable and interpretable strategy for assessing cardiomyocyte maturation, disease remodeling, and therapeutic response using AI models.

### Limitations of The Study

Cardiomyocyte function is inherently time-dependent, involving cyclic sarcomere shortening, relaxation kinetics, and contraction–relaxation coupling. However, in this study, contractile function was represented as a static intensity-based heatmap, capturing the spatial magnitude and distribution of contraction but not its temporal dynamics. Consistent with this limitation, AI-derived metrics showed weak correlation with beating dynamics such as beat rate and contraction duration, indicating that static morphology alone cannot fully encode temporal behavior or excitation–contraction coupling. Future work will incorporate temporal modeling into the framework. Integration of time-series imaging with recurrent or temporal convolutional networks could enable prediction of contraction kinetics. Emerging diffusion-based generative models^41,42^ may further allow reconstruction of dynamic “virtual beating” sequences from static cellular morphology.

In addition, the current framework relies on image-to-function translation learned from a relatively limited dataset, which may constrain generalizability across different cell lines, differentiation protocols, or imaging conditions. Although data augmentation using synthetic hiPSC-CMs improved model robustness, a potential domain gap between synthetic and experimentally derived cells remains. While quality control measures were applied, synthetic data may not fully capture subtle biological variability or rare phenotypes, which could introduce bias in model training. Similarly, ROI-based analyses and thresholding strategies, while systematic, may introduce sensitivity to parameter selection and may not fully represent continuous biological gradients. Improving cardiomyocyte maturity through optimized culture strategies—such as enhanced mechanical conditioning and metabolic maturation—may strengthen structure–function fidelity and improve predictive accuracy. Finally, extending the model from single cells to three-dimensional cardiac organoids and engineered heart tissues will be essential to capture multiscale organization and increase translational relevance. Together, these advancements will enable a more comprehensive AI framework capable of modeling the integrated spatial and temporal dynamics of cardiac function.

## METHODS

### hiPSC-CM Differentiation and Purification

The ACTN2-GFP hiPSC line (Cell Line ID: AICS-0075 cl.85) was engineered by the Allen Institute and obtained from the Coriell Institute for Medical Research. hiPSCs were cultured on Geltrex-coated plates (Cat# A1413302; Thermo Fisher Scientific) and maintained in Essential 8 (E8) medium (Cat# A1517001; Thermo Fisher Scientific). Cells were passaged upon reaching ~90% confluency. During the first 24 hours after passaging, the medium was supplemented with 10 μM ROCK inhibitor (Y-27632; STEMCELL Technologies, Cat# 72308) to enhance cell survival.

Cardiomyocyte differentiation was initiated using a small-molecule WNT modulation protocol^43^. On day 0, hiPSCs were treated with 10 µM GSK3 inhibitor (CHIR99021; STEMCELL Technologies) in RPMI 1640 medium supplemented with B27 minus insulin (B27–I) to induce mesoderm formation for 24 hours. On day 1, the medium was replaced with fresh RPMI/B27–I for recovery. On days 3–5, cells were treated with 5 µM WNT inhibitor (IWP4; STEMCELL Technologies) in RPMI/B27–I to promote cardiac specification. From day 6 to day 12, cells were maintained in RPMI supplemented with B27 complete (B27+C) medium. Differentiated hiPSC-CMs were dissociated using the STEMdiff Cardiomyocyte Dissociation Kit (STEMCELL Technologies). Metabolic purification was performed using a lactate selection medium consisting of glucose-free DMEM (Thermo Fisher Scientific), non-essential amino acids (NEAA; Thermo Fisher Scientific), GlutaMAX (Thermo Fisher Scientific), and 4 mM sodium L-lactate (Sigma-Aldrich). Purified cardiomyocytes were subsequently maintained in RPMI/B27+C medium, with media changes every two days.

### hiPSC-CM Micropatterning

A silicon wafer was coated with SU-8 50 and spun using a photoresist spinner to obtain a thickness of 25 µm. The coated wafer was pre-baked at 65 °C for 3 min, followed by a soft bake at 95 °C for 7 min, then allowed to cool to room temperature on a flat surface. The photomask was placed on top of the SU-8–coated wafer and exposed to UV light. The exposed wafer then underwent a post-exposure bake at 65 °C for 1 min and 95 °C for 3 min, at which point the patterns became visible. Uncrosslinked SU-8 was removed by immersing the wafer in SU-8 Developer for 4 min, followed by two rinses with isopropanol (IPA) and drying with nitrogen.

Crosslinked PDMS elastomers was prepared from a Sylgard-184 kit by mixing with the base and curing agent at a 10:1 ratio. The mixture was poured onto the patterned silicon wafer, then cured at 60 °C overnight to ensure complete crosslinking and replicating the patterns. After curing, the PDMS was carefully peeled from the wafer and cut into small stamps for microcontact printing. PDMS stamps were sterilized under UV light and incubated with rhodamine labeled fibronectin (FN) (100 µg/mL) at 37°C for 1 h to allow protein adsorption onto the stamp surface. The stamps and glass substrates were transferred to an environmental control room with constant 50% relative humidity. The humidified environment improves FN transfer from the stamps to the substrates. PDMS stamp was pressed onto a glass substrate under a 20 g weight for 2 min. While preparing cells for seeding, glass substrates were passivated with 0.4% Pluronic F-127 for 1 h. Excessive Pluronic will be removed by three PBS washes. hiPSC-CMs were dissociated into single cells and seeded at 2 × 10^4^ cells/mL onto fibronectin-patterned glass substrates for 4 days before microscopy.

### Video Recording and Contraction Analysis

Brightfield and fluorescence images were acquired using a Nikon Eclipse Ti inverted microscope equipped with a Zyla 4.2 PLUS sCMOS camera. Brightfield videos of spontaneously beating hiPSC-CMs were recorded at 50 frames per second for 10 seconds. Immediately following each brightfield acquisition, a corresponding fluorescent image of sarcomere organization (ACTN2-GFP) from the same cell was captured, ensuring paired datasets of cellular morphology, sarcomere structure, and contractile motion.

Contractile motion was quantified from brightfield videos using an open-source, video-based block-matching motion analysis algorithm^9^. Raw videos were cropped to 575 pixels × 575 pixels regions centered on individual cells to standardize input dimensions. For each frame, displacement vectors were computed to generate motion-field matrices describing local pixel movement (Supplemental Figure 5). Displacement vectors were then aggregated across all frames to produce a contraction heatmap representing the mean contraction intensity per pixel. These heatmaps provided a quantitative spatial map of the magnitude and distribution of contractile motion for each individual hiPSC-CM. In addition to generating spatial contraction heatmaps, the motion-tracking analysis also extracted quantitative beating parameters, including beat rate, maximal contraction velocity, maximal relaxation velocity, and the contraction–relaxation interval.

### U-Net-GAN Model for Contraction Prediction

Our AI model builds upon the U-Net autoencoder architecture^44^, which encodes the input image through sequential convolutional layers into a compact latent representation and reconstructs it via symmetric transposed convolutions. Skip connections between corresponding encoder and decoder layers preserve fine spatial details and high-frequency features critical for accurate image translation. The generator consists of a 7-layer encoder–decoder network using 4×4 convolutional kernels with stride 2, batch normalization, and leaky ReLU activations in the encoder. The decoder comprises 7 transposed convolutional layers with skip connections linking each encoder– decoder pair to retain spatial information. A final 1×1 convolution followed by a Tanh activation produces a 2D RGB heatmap representing predicted contraction intensity across the cell.

To enhance output realism and structural fidelity, we extended the U-Net generator within a pix2pix generative adversarial network (GAN) framework^45^. In this configuration, the U-Net generator is trained jointly with a 70×70 PatchGAN discriminator. Rather than evaluating images globally, the PatchGAN discriminator operates on local image patches, serving as a conditional classifier that enforces high-resolution, context-aware image translation. The generator loss function (ℒ_*G*_) integrates three components: adversarial loss (ℒ_*GAN*_) to encourage realistic heatmap synthesis, reconstruction loss (ℒ_*L*1_) to enforce pixel-wise fidelity to ground truth, and a structural similarity term (SSIM) to preserve perceptual and spatial consistency. We adopted initial weights of *λ*_*GAN*_ = 1.0, *λ*_*L*1_ = 100.0, and *λ*_*SSIM*_ = 1.0 (Isola et al. (2017)).

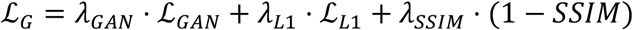

The structural similarity index (SSIM) quantifies image similarity as the product of three components: luminance (l), which compares mean intensity; contrast (c), which reflects differences in standard deviation; and structure (s), which captures spatial correlation between images. SSIM was computed using the scikit-image library in Python^46^ to enable efficient evaluation during model training and testing. Higher SSIM values indicate greater structural agreement between the predicted contraction heatmaps and the corresponding ground-truth images.

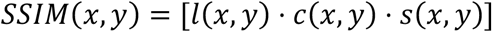

In addition to SSIM, we implemented the Learned Perceptual Image Patch Similarity (LPIPS) metric to further quantify the accuracy of contraction predictions. LPIPS is a perceptual image similarity metric that compares deep feature representations between two images rather than relying solely on pixel-wise differences. Specifically, we used the AlexNet-based variant (LPIPS-alex), which extracts multi-layer features using a pretrained AlexNet backbone and applies learned linear weights to compute a single scalar similarity score. Lower LPIPS values indicate higher perceptual similarity between the predicted and ground-truth images^47^.

All models were implemented in Python (v3.12) using TensorFlow 2.11 with the Keras API and trained on Google Colab Pro GPUs. Input images were resized to 256 × 256 pixels to match the U-Net architecture requirements. Batch size was determined based on available GPU memory, and training duration was monitored every 100 epochs to assess convergence and prevent overfitting. Optimization was performed using the Adam optimizer (β_1_ = 0.5, β_2_ = 0.999) with a learning rate of 2 × 10^−4^. Data augmentation, including random horizontal flipping, contrast adjustment, and gamma shifting, was applied to improve model generalizability. Model checkpoints were saved every 10 epochs, and the final model was selected based on performance on the testing set using SSIM and LPIPS-alex metrics. All testing data were strictly withheld during training and were used exclusively for independent evaluation against paired image–heatmap ground-truth datasets.

A baseline U-Net model and a U-Net-GAN pix2pix model are trained and compared. The standalone U-Net was trained by optimizing the generator to minimize a combination of L1 loss and SSIM, emphasizing pixel-level accuracy and structural similarity. In contrast, the pix2pix model jointly trained both the generator and discriminator. The L1 loss primarily enforced pixel-wise accuracy, while the SSIM loss preserved structural consistency between predicted and ground-truth heatmaps. Training dynamics for both generator and discriminator losses are shown in Supplemental Figure 6.

### StyleGAN2 Model for Synthetic hiPSC-CM generation

A StyleGAN2-ADA framework was employed to generate synthetic overlay images of hiPSC-CMs paired with corresponding contraction heatmaps. The model was adapted from NVIDIA’s official PyTorch implementation and trained in a Google Colab T4 GPU environment for 200 kimg^48^. Training followed standard StyleGAN2 hyperparameters, with adaptive discriminator augmentation (ADA) enabled to stabilize training on the relatively small dataset. ADA dynamically adjusts augmentation probability to prevent discriminator overfitting while preserving biologically relevant variability. To learn the joint distribution between cellular structure and contractile function, paired training images were constructed by concatenating brightfield morphology and contraction heatmaps into a single composite image. After training, synthetic paired samples were generated and subjected to quantitative quality control to ensure fidelity of structure–function correspondence. After training, synthetic paired images were generated and manually curated to remove hallucinations. Only high-quality samples preserving realistic alignment between cellular morphology and contraction patterns were retained.

From an initial pool of ~120 synthetic samples, 81 high-quality paired images passed filtering criteria and were retained. These were combined with the original experimental dataset (n = 296 cells), resulting in an augmented training dataset of n = 377 samples. The synthetic cell–heatmap pairs were then split into individual channels, cropped, and preprocessed to match the input format of the U-Net–GAN model. Unlike deterministic mapping approaches that constrain variability, the StyleGAN2 framework learns a latent distribution of hiPSC-CM features, enabling the generation of diverse yet realistic synthetic samples that expand the diversity of cell morphology and sarcomere organization. This allows exploration of feature spaces that may be underrepresented in experimental datasets while maintaining biological consistency. All preprocessing steps, including pairing, filtering, and formatting, were implemented within the StyleGAN pipeline and are available in the shared code repository.

### Automated Region-of-Interests (ROIs) Selection

Two distinct ROI categories were defined based on ground-truth and AI-predicted heatmaps. An initial global ROI was first generated from the predicted heatmap using Otsu thresholding to delineate the valid cell region. For contraction-based segmentation, ground-truth heatmaps were used to identify high- and low-contraction ROIs based on relative color intensity. High-contraction regions were defined using thresholds on the RGB channels to capture the upper intensity spectrum of the colormap: pixels with red channel intensity > 0.5, or red > 0.7 combined with green > 0.9 (capturing orange–yellow regions), were classified as high-contraction ROIs. Low-contraction regions were defined as pixels within the heatmap mask exhibiting blue channel intensity > 0.7, excluding areas already classified as high contraction. This color-based thresholding strategy leverages the predefined colormap mapping between intensity and RGB values, providing a consistent and computationally efficient proxy for segmenting contraction magnitude. For error-based segmentation, AI-predicted heatmaps were compared with ground-truth heatmaps to compute pixel-wise reconstruction error. High-error ROIs were defined as pixels within the upper 55th percentile of the error distribution, while the remaining pixels within the cell mask were classified as low-error ROIs. All ROIs were subsequently filtered using a minimum area threshold of 100 pixels to ensure sufficient spatial coverage for sarcomere visualization and downstream 2D FFT analysis. Prior to FFT computation, a 20-pixel background subtraction was applied to each ROI to reduce edge artifacts and improve spectral resolution.

### Sarcomere Analysis and Two-Dimensional Fast Fourier Transform (2D FFT)

The computational package integrates automated batch-analysis of sarcomere organization. Fluorescent ACTN2-GFP images of hiPSC-CMs were processed using Otsu thresholding to segment sarcomere structures. To quantify higher-order sarcomere organization, we applied 2D FFT analysis^24^. For each cell or sarcomere ROI, a 2D FFT was computed using the SciPy FFT module to transform spatial intensity patterns into the frequency domain (Supplemental Figure 7). The magnitude spectrum was normalized to its maximum amplitude and centered at zero frequency. A radial average of the FFT magnitude was then calculated by averaging spectral intensities as a function of radial distance from the center, yielding a 1D frequency profile that captures global periodicity. From this radial profile, the fundamental frequency peak was identified to estimate average sarcomere length (inverse of frequency). The full width at half maximum (FWHM) of the peak was extracted as a measure of sarcomere regularity, with broader peaks indicating increased inhomogeneity in sarcomere length. In addition, the dominant orientation angle of the FFT magnitude spectrum was computed to quantify sarcomere orientation within each cell. In addition, sarcomere density was calculated as the fraction of fluorescent area relative to the total cell or ROI area.

### Cell Morphology Analysis

The computational package integrates automated batch analysis of cell morphology using functions from scikit-image^46^. Grey and binary masks were generated from brightfield images using Otsu thresholding, followed by min– max normalization to standardize intensity. Cell perimeter was quantified using both standard perimeter and Crofton perimeter estimators implemented in scikit-image. Circularity was calculated as 4π *×* area*/*perimeter^)^, providing a measure of shape compactness. Additional morphological features, including major and minor axis lengths and aspect ratio, were extracted from region properties derived from the binary cell mask.

### Two-Feature Logistic Regression Classification

A two-feature logistic regression classification framework was applied to six pairwise combinations of four sarcomere descriptors: length, regularity, orientation, and density. Binary outcome labels were assigned by median split of SSIM values across the n = 30 test cells, with cells at or above the median SSIM (0.842) designated as the high-predictability class (positive label = 1) and cells below the median as the low-predictability class (label = 0). For each feature pair, input features were z-score normalized (zero mean, unit variance) using StandardScaler from scikit-learn. A logistic regression classifier with L2 regularization (C = 1.0, max_iter = 1000) was fitted using the LogisticRegression implementation from scikit-learn (Python v3.12). Model performance was estimated using leave-one-out cross-validation (LOO-CV) and quantified using the area under the receiver operating characteristic curve (AUC-ROC). AUC values ≥ 0.75 were interpreted as indicative of good discriminative ability. For visualization, a logistic regression model was fitted to the full dataset to generate decision boundary plots with a 0.50 probability contour in the two-dimensional feature space. Background probability fields were rendered as color-filled contour maps (green = high predicted SSIM probability, red = low). Classification accuracy, sensitivity, and specificity were computed from the full-data fitted model predictions. All analyses were conducted in Python v3.12 using scikit-learn, NumPy, SciPy, and Matplotlib.

### Statistical Analysis

All preprocessing and quantitative analyses were performed in Python (v3.12). Image handling and visualization were conducted using OpenCV, while NumPy, Matplotlib, and Seaborn were used for numerical computation and data visualization. Inferential statistical analyses were performed using SciPy and Statsmodels. Two-tailed Student’s t-tests were applied for pairwise comparisons, and one-way analysis of variance (ANOVA) followed by Tukey’s post hoc test was used for multiple-group comparisons. Statistical significance was defined as p < 0.05 (*), < 0.01 (**), < 0.001 (***), and < 0.0001 (****).

## Supplemental Information

Supplemental information can be found online.

## Acknowledgement

This work was supported by the NIH [R01HD101130, R21HD11458, R15HD108720], the NSF [2130192 and 1943798].

## Author Contributions

A.K. and Z.M. conceived the study. A.K. performed all the experiments and collected raw data. C. W. helped with cell differentiation. X.W. provided insights on cardiomyocyte structure-function relationship. H.Y. and Z.Q. provided insights on AI models and integration with biological systems. Z.M. supervised the project development. A.K. and Z.M. wrote the manuscript with discussion and improvements from all authors. Z.M. funded the study.

## Resource Availability

The code for generative AI for contraction prediction of the single cardiomyocytes is available at https://github.com/ankowalc/SingleCell_hiPSC_CM_Analysis

Further information and requests for complete dataset of micropatterned hiPSC-CMs should be directed to and will be fulfilled by the lead contact, Zhen Ma.

