## Supplemental Figures for "A Generative AI Framework to Predict Cardiomyocyte Contraction Function from Single Static Images"

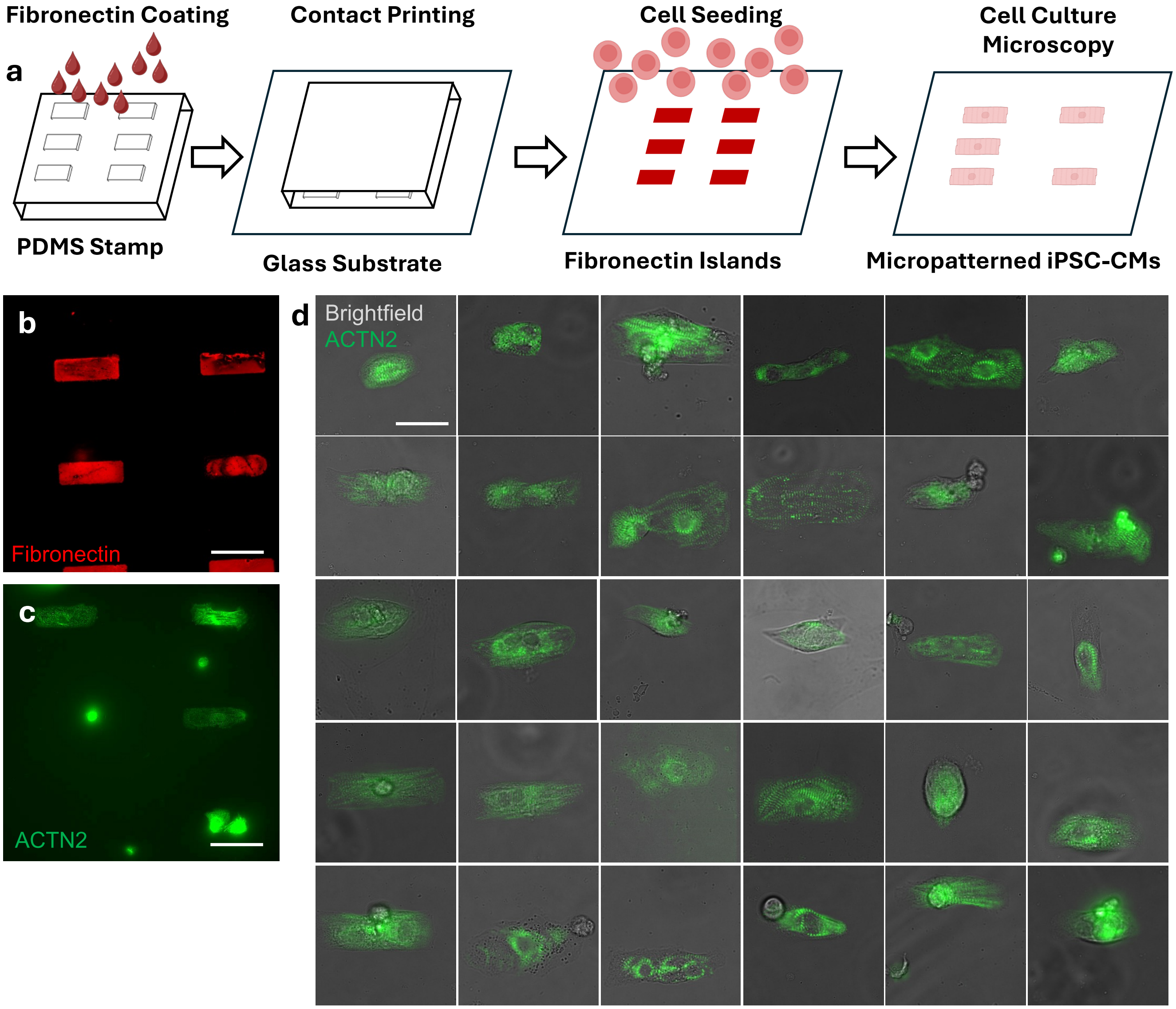


**Supplemental Figure 1. Micropatterning of hiPSC-CMs.** (a) Schematic of the microcontact printing process and hiPSC-CM micropatterning workflow. (b) Micropatterned array of fibronectin islands. (c) hiPSC-CMs cultured on fibronectin-patterned substrates. (d) Overlay images of cell morphology (brightfield) and sarcomere structure (fluorescence) from all 30 hiPSC-CMs in the testing dataset.

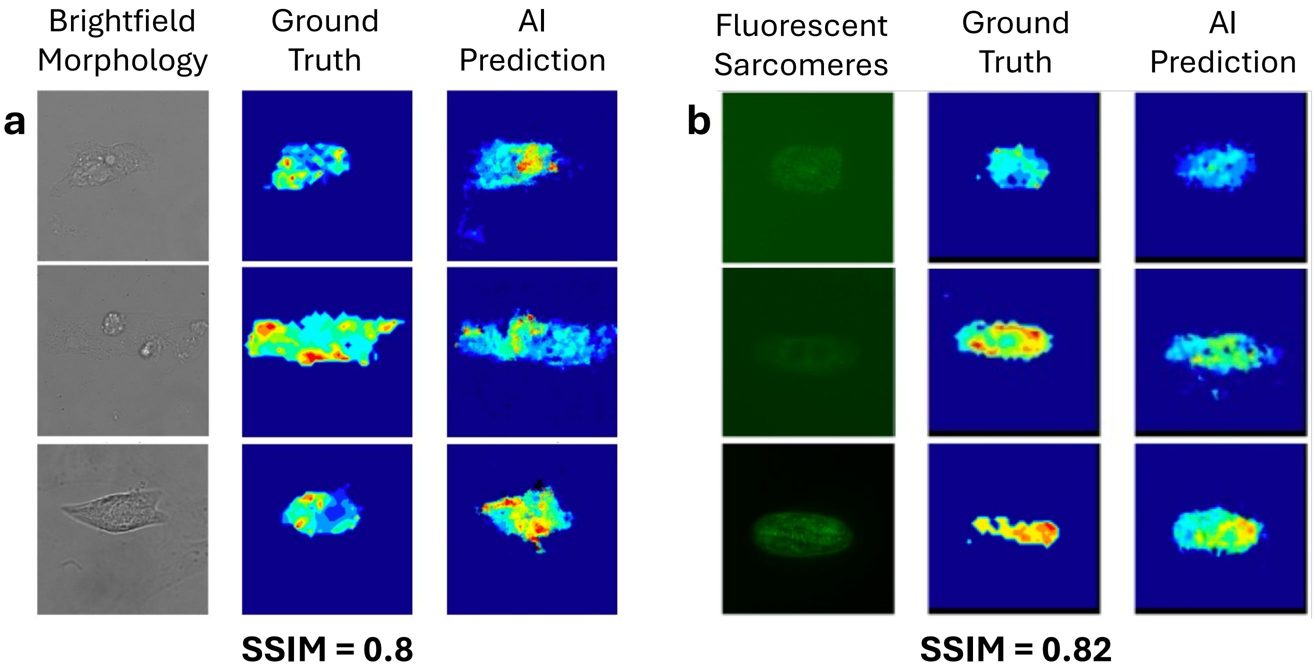


**Supplemental Figure 2. Predict contraction heatmaps from static images of cell morphology and sarcomere structure.** (a) The U-Net–GAN translates brightfield cell morphology images into contraction heatmaps with an SSIM of 0.80, (b) while fluorescent sarcomere structure images yield improved predictions with an SSIM of 0.82.

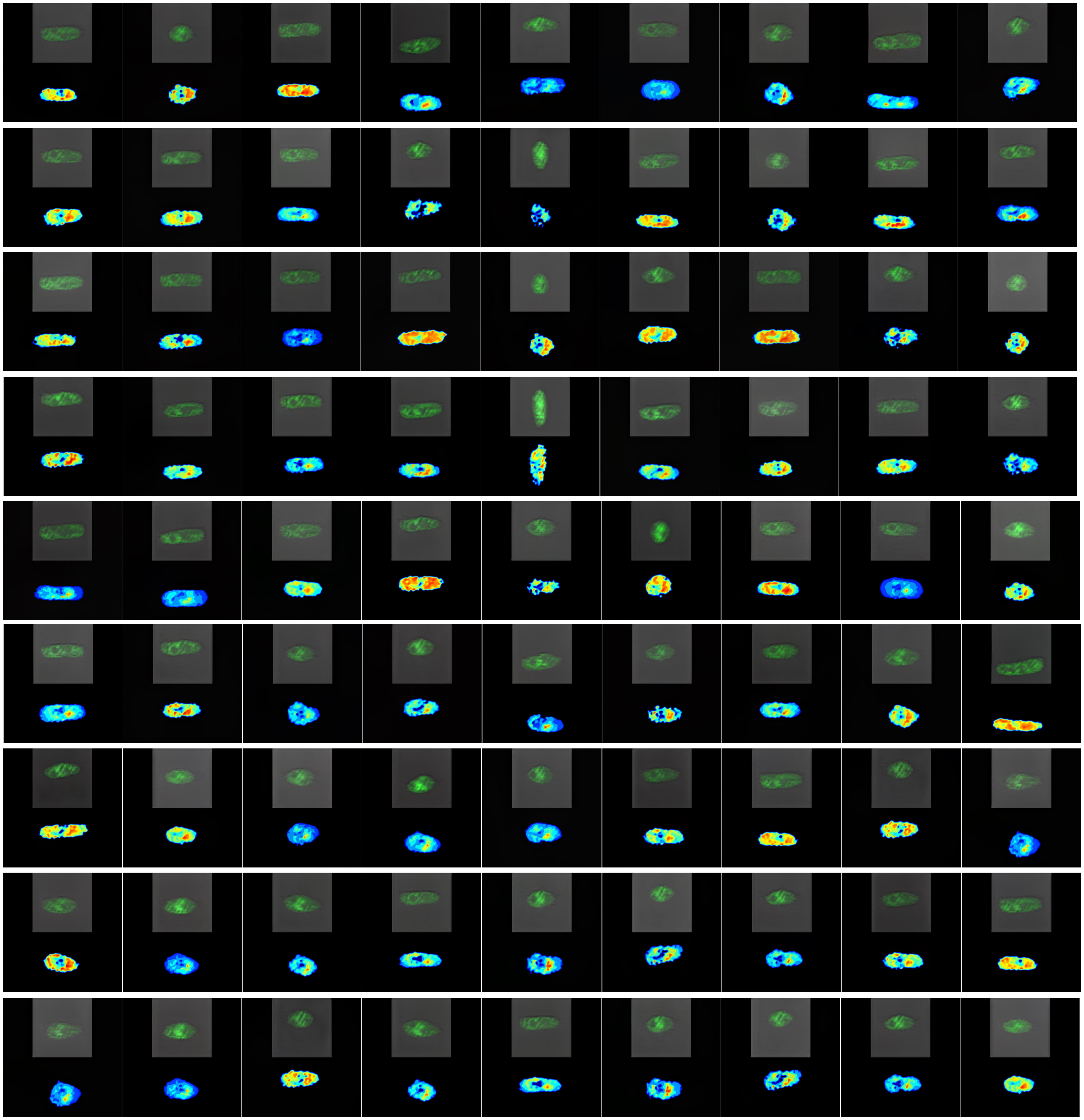

**Supplemental Figure 3. Synthetic hiPSC-CMs generated by the StyleGAN2 model.** All 81 synthetic hiPSC-CM images paired with corresponding contraction heatmaps used for training the U-Net–GAN model.

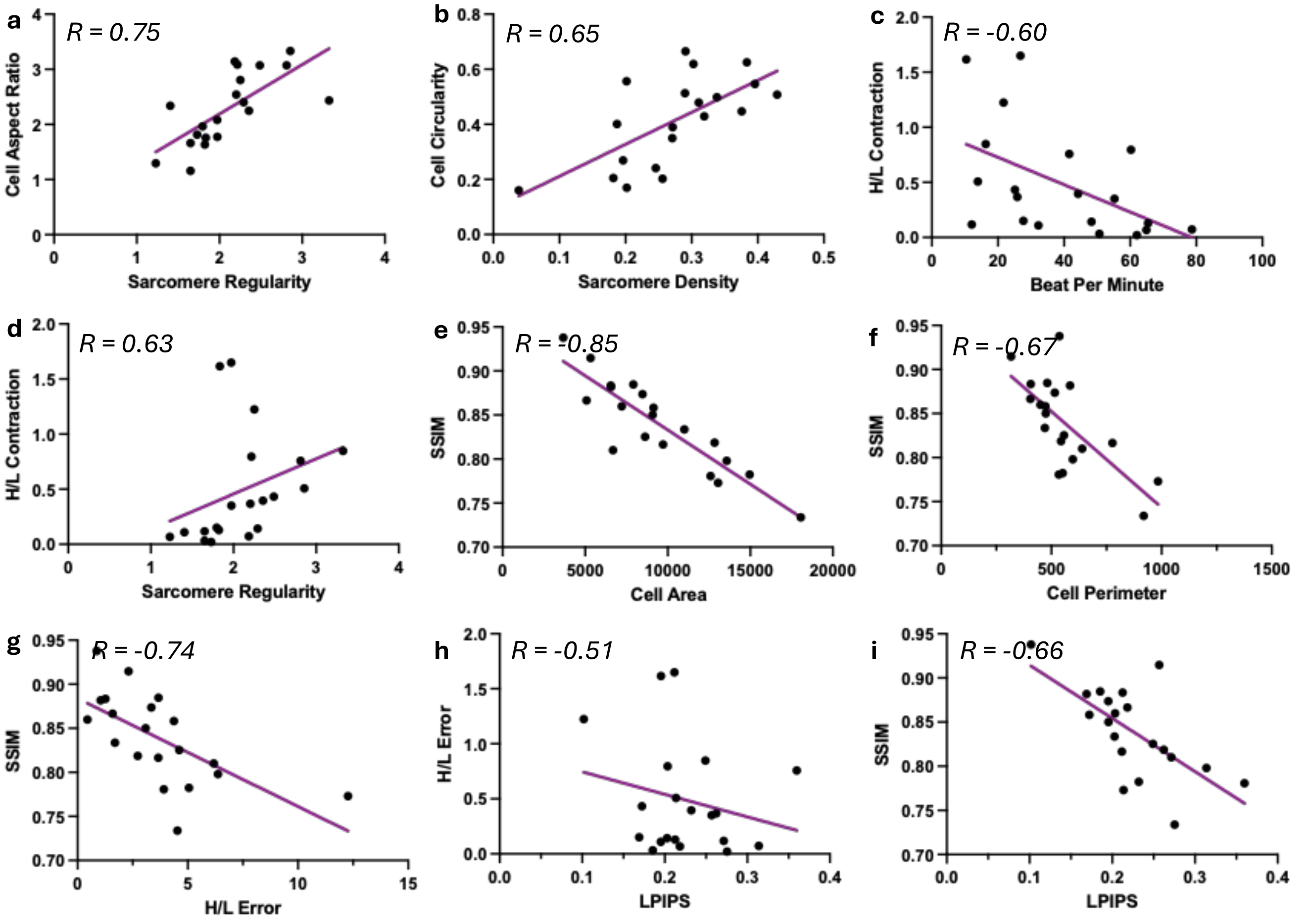


**Supplemental Figure 4. Linear regression analysis of cell morphology, sarcomere structure, contractile function, and AI metrics.** Selected linear relationships with correlation coefficients greater than 0.5 are shown: (a) cell aspect ratio – sarcomere regularity; (b) cell circularity – sarcomere density; (c) H/L contraction – beat per minute; (d) H/L contraction – sarcomere regularity, (e) SSIM – cell area; (f) SSIM – cell perimeter; (g) SSIM – H/L error; (h) LPIPS – H/L error; and (i) SSIM – LPIPS.

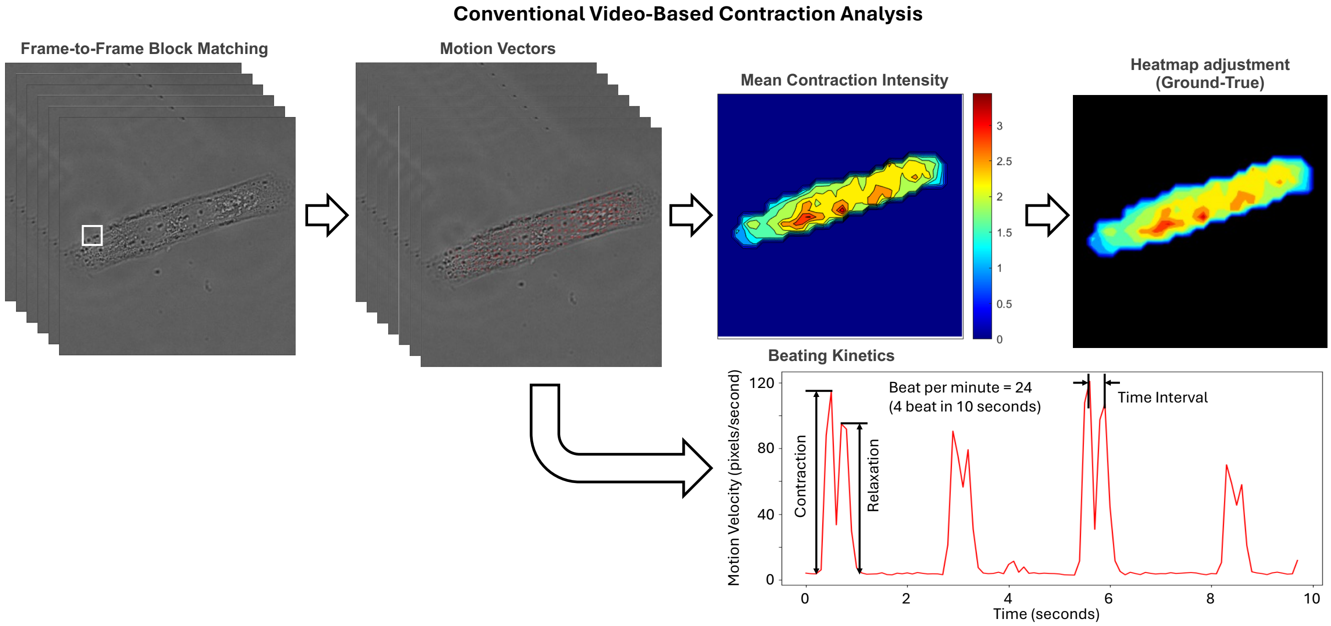


**Supplemental Figure 5. Conventional video-based contraction analysis.** Brightfield image stacks of beating hiPSC-CMs were analyzed using a block-matching algorithm to generate mean contraction intensity heatmaps and extract quantitative beating kinetics for contraction feature analysis.

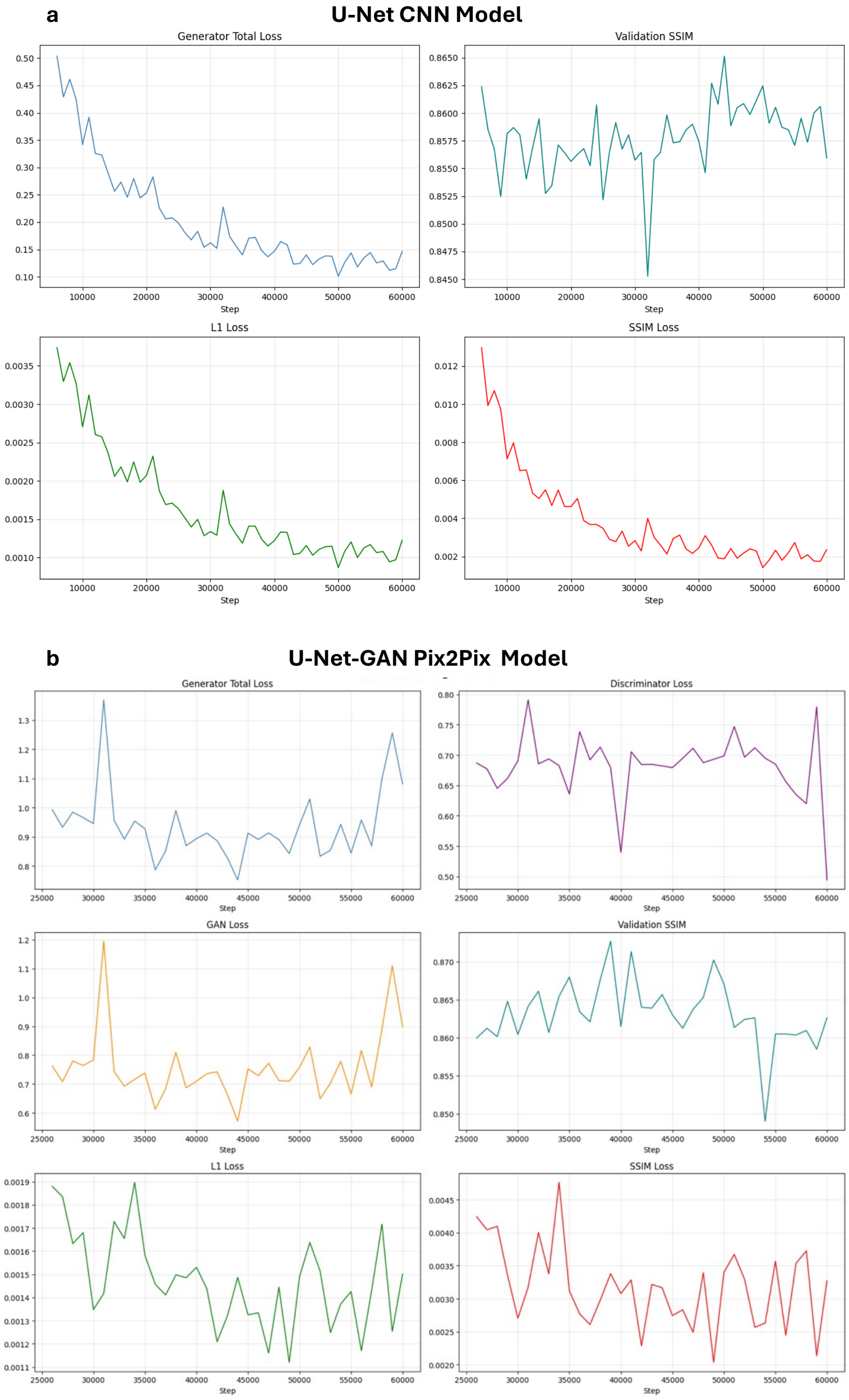


**Supplemental Figure 6. Representative Training Loss Curves.** (a) Training curves for the baseline U-Net model. The generator total loss, L1 loss, and SSIM loss show stable and monotonic convergence, indicating effective optimization. (b) Training curves for the U-Net–GAN (pix2pix) model. Generator and discriminator losses exhibit characteristic oscillatory behavior due to adversarial training dynamics. The GAN loss reflects the competitive interaction between generator and discriminator, while L1 and SSIM losses remain stable.

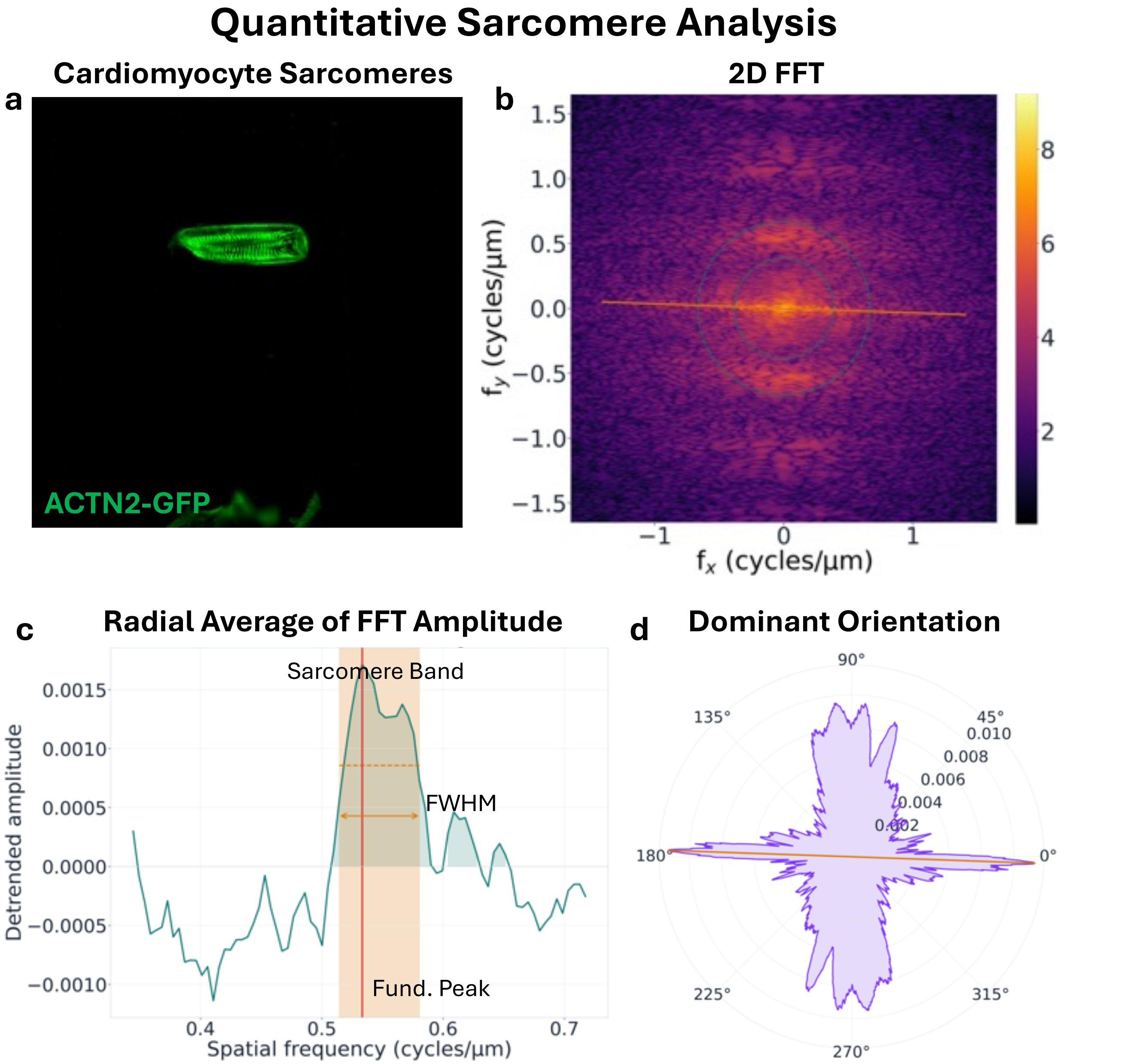


**Supplemental Figure 7. Quantitative sarcomere analysis using 2D FFT.** (a) Representative fluorescence image of sarcomere structure in a micropatterned hiPSC-CM. (b) Corresponding 2D FFT magnitude spectrum computed from the sarcomere image. (c) Radially averaged 1D FFT profile was detrended to identify the dominant frequency peak for quantifying sarcomere spacing and regularity. (d) Dominant orientation angle extracted from the FFT magnitude spectrum to quantify sarcomere alignment.
